# Spatio-temporal variation of bacterial community structure in two intertidal sediment types of Jiaozhou Bay

**DOI:** 10.1101/2023.05.24.542048

**Authors:** Xuechao Chen, Xinran Zhang, Hao Yu, Meiaoxue Han, Jianhua Sun, Gang Liu, Yan Ji, Chuan Zhai, Liyan Zhu, Hongbing Shao, Yantao Liang, Andrew McMinn, Min Wang

**Author notes:** Corresponding author, E-mail addresses (M. Wang), (A. McMinn), (Y. Liang). These authors contributed equally to this work (X. Chen, X. Zhang and H. Yu).

## Abstract

The intertidal sediment environment is dynamic and the biofilm bacterial community within it must constantly adjust, but an understanding of the differences in the biofilm bacterial community within sediments of different types is still relatively limited. In this study, the structure of the bacterial community in Jiaozhou Bay sediment biofilms are described using high-throughput 16S rRNA gene sequencing and the effects of temporal change and different sediment environment types are discussed. The Shannon index was significantly higher in sandy samples than in muddy samples. The co-occurrence network was tighter and more species were involved in community building in sandy samples. The principal coordinates analysis identified a significant separation between different sediment types and between stations (LiCun estuary, LC and ZhanQiao Pier, ZQ). Proteobacteria, which had a relative abundance of approximately 50% at all phylum levels, was significantly more abundant at ZQ, while Campilobacterota and Firmicutes were significantly more abundant at LC. The relative abundances of Bacteroidetes, Campilobacterota, Firmicutes, and Chloroflexi were significantly higher in the muddy samples, while Actinobacteria and Proteobacteria were higher in the sandy samples. There were different phylum-level biomarkers between sediment types at different stations. There were also different patterns of functional enrichment in biogeochemical cycles between sediment types and stations with the former having more gene families that differed significantly, highlighting their greater role in determining bacterial function. The RDA results, where each month’s samples were concentrated individually, showed reduced variation between months when the amplicon sequence variant was replaced by KEGG orthologs, presumably the temporal change had an impact on shaping the intertidal sediment bacterial community, although this was less clear at the gene family level. Random forest prediction yielded a combination of 43 family-level features that responded well to temporal change, reflecting the influence of temporal change on sediment biofilm bacteria.

**Highlights:**

- Sandy sediments have more bacterial species involved in community building.
- Different substrates from different stations have their own phylum biomarkers.
- Substrates have a greater influence on shaping bacterial function.
- Temporal changes have a greater shaping power on bacteria than on gene families.

**Graphic abstract:** 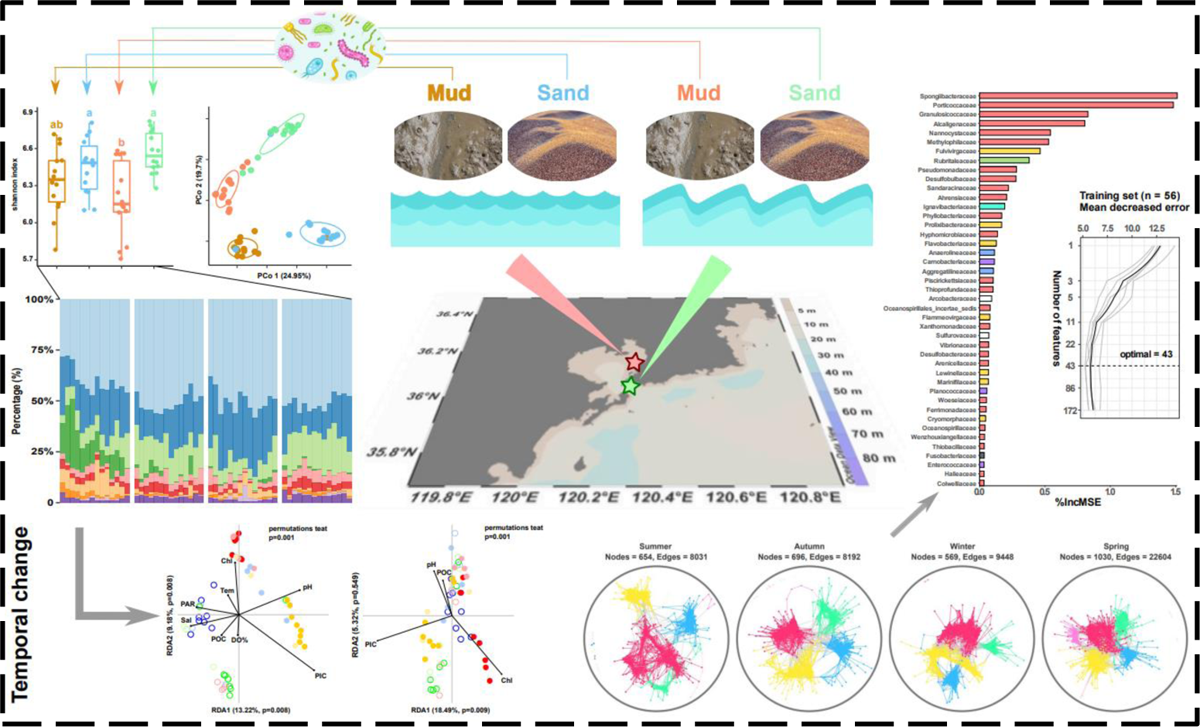

## Introduction

The intertidal zone is a transition zone between land and sea where the sediment environment is constantly changing under the combined influence of freshwater, tidal and anthropogenic factors. Studies have shown that the sediment surface has high microbial activity, which is sensitive to environmental changes and can be an indicator of ecological health (Duarte et al., 2012; Kallmeyer et al., 2012; Suh et al., 2015; Yi et al., 2020). In addition to microorganisms on the surface of intertidal sediments that adhere and aggregate into marine biofilms, marine biofilms are also widespread on the surface of other biotic or abiotic substrates immersed in seawater and play an important role in biogeochemical cycles (Wahl et al., 2012; Dang and Lovell, 2016; Liang et al., 2019).

Bacteria are the pioneer and most abundant taxa in biofilms, and their diversity and aggregation state are significantly influenced by water quality, salinity and hydrodynamics (Rickard et al., 2004; Nocker et al., 2007; Qian et al., 2007; Guo et al., 2017). Rapid fluctuations in intertidal environmental conditions (e.g., pH, salinity, ionic strength, currents and contaminants) result in a metabolically plastic and genetically diverse community of bacteria. Understanding how changes in the distribution and function of these communities and the extent to which anthropogenic and natural factors influence them is essential for assessing ecological impacts (Lv et al., 2016; Wei et al., 2018; Avila-Jimenez et al., 2020; Ge et al., 2021). With the development and maturity of sequencing technology, in particular using high-throughput 16S rRNA gene sequencing technology, it has become possible to study community composition at a lower cost, with greater accuracy and efficiency.

Tidal, current and invertebrate disturbance have different effects on the erosion of the surface layer of intertidal sediments with different grain sizes (Grabowski et al., 2011). Sandy sediments have high permeability but low interparticle cohesion, whereas muddy sediments have smaller particle sizes, higher cohesion and increased stability, so surface migration differs between the two sediment types. In addition, sediment mobility is also influenced by the biofilms themselves (Paterson, 1989; Whitehouse, 2000; Wyness et al., 2019). However, an understanding of the structural differences in intertidal biofilm bacterial communities in different substrate types is still relatively limited. Therefore, it is hypothesised that intertidal muddy and sandy sediments shape bacterial communities differently. In this study, Jiaozhou Bay, a temperate bay with different sediment types, was chosen as a study model.

Jiaozhou Bay (Shandong, China) is a semi-enclosed bay with a relatively narrow mouth (2.5 km) that restricts water exchange to the Yellow Sea and limits its capacity to self-purify (Dai et al., 2007; Sun et al., 2021). It’s semi-enclosed configuration results in low wave energy. It has a temperate monsoon climate, semi-diurnal tides (mean tidal range 2.8 m) and high current velocities (up to 201 cm/s) at the mouth of the bay (Lyu et al., 2010; Liu et al., 2014; Shang et al., 2018). The Licun River, which flows into the bay, has a population of more than one million people within its catchment. A large sewage treatment plant discharges into the Licun River (2.46 × 10^5^ t/d). A eddy in the estuary attenuates the movement of material out of the bay (Wang et al., 2022; Zhang et al., 2017). Previous studies have investigated the diversity of culturable bacteria (Wang et al., 2016), the diversity of anaerobic bacteria (Wu et al., 2019), the *nirS*-type denitrifying bacterial community (Liu et al., 2020), the late winter/early spring bacterial community in the estuarine zone (Ge et al., 2021), and the spatial distribution of bacteria in autumn (Sun et al., 2021) in surface sediments of Jiaozhou Bay, but investigations of bacterial community structure between seasons and substrates are need to be complemented.

In a monthly sampling program for almost a year, the bacterial community structure of the intertidal sediments in the bay and at the mouth of the bay is described using high-throughput 16S rRNA gene sequencing methods. The aim is to identify potential patterns in the composition of bacterial communities in the sediment biofilms, and the effects of temporal changes and sediment environment on them. The results will improve our understanding of the endemic structure of bacterial communities shaped by different substrate sediments in the intertidal zone and provide the empirical support necessary for ecological conservation.

## Materials and Methods

### Sampling design

The sampling sites were at two locations in Jiaozhou Bay - the south side of the LiCun estuary (LC, 36.15 N 120.34 E) and a beach on the east side of the ZhanQiao Pier (ZQ, 36.06 N 120.31 E), inside and outside the mouth of the bay respectively, a linear separation of approximately 10.5 km. Samples were collected monthly over a period of 326 days (Fig. 1). Sampling preferentially took place when low tides (<90 cm) occurred close to midday. Surface samples (0.5 cm) were collected from different sediment types (muddy samples, sandy samples). The samples from each site were collected close to the water’s edge (where the samples were only exposed to air for a short time, resulting in less water loss and less additional stress), divided into collection tubes for cold storage at −80 °C in the laboratory. Water temperature (Tem), dissolved oxygen air saturation (DO%), dissolved oxygen (DO), total dissolved solids (TDS), salinity (Sal), pH of the seawater adjacent to the sediment were measured *in situ* using a YSI water quality analyzer. Photosynthetically active radiation (PAR), chlorophyll (Chl), particulate inorganic carbon (calcite concentration, PIC), particulate organic carbon (POC) were extracted from MODIS-Aqua remote sensing data, and if the extracted values were NA, then extracted from 8-day average, monthly average, etc.

**Fig. 1.**
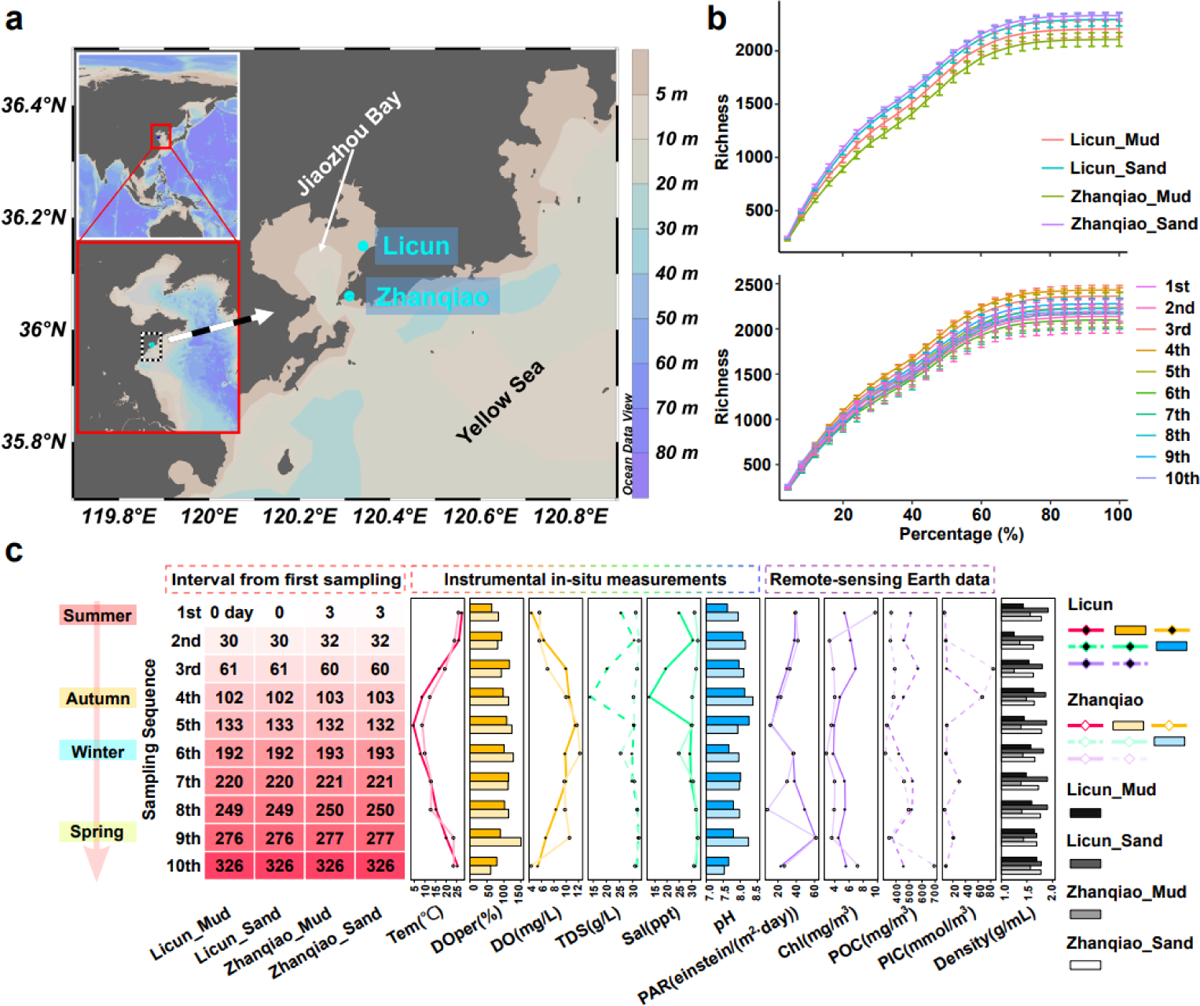
Sampling design. a) Geographical information on the stations b) Richness dilution curve c) Sampling time and variation of environmental factors (Tem: water temperature Doper: dissolved oxygen air saturation DO: dissolved oxygen TDS: total dissolved solids Sal: salinity PAR: photosynthetically active radiation PIC: particulate inorganic carbon POC: particulate organic carbon).

### High throughput sequencing

Samples for sequencing were collected each month, with the August (1^st^ sample month), November (4^th^), February (6^th^) and May (9^th^) being characteristic months of each local season; an additional parallel sample was also sequenced. DNA was extracted from the genomic DNA of the microbial community according to the instructions of the E.Z.N.A.® Soil DNA Kit (Omega Bio-tek, Norcross, GA, USA). 1% agarose gel electrophoresis was used to check DNA quality and NanoDrop 2000 (Thermo Scientific, USA) was used to determine DNA concentration and purity. PCR amplification of the V3-V4 variable region of the 16S rRNA gene was performed using upstream primer 338F (5’-ACTCCTACGGGAGGCAGCAG-3’) and downstream primer 806R (5’-GGACTACHVGGGTWTCTAAT-3’) with barcode sequences (Liu et al., 2016), the PCR reaction system was: 5×FastPfu buffer 4 μL, 2.5 mM dNTPs 2 μL, each primer (5 μM) 0.8 μL, Fast Pfu polymerase 0.4 μL, bovine serum albumin (BSA) 0.2 μL, template DNA 10 ng, and ddH2O to a final volume of 20 µL. Amplification procedure: initial denaturation at 95°C for 3 min, 27 cycles (denaturation at 95°C for 30 s, annealing at 55°C for 30 s, extension at 72°C for 45 s), followed by a single extension at 72°C for 10 min (ABI GeneAmp® 9700). Three replicates of each sample were run. PCR products from the same sample were mixed and recovered on a 2% agarose gel, purified using the AxyPrep DNA Gel Extraction Kit (Axygen Biosciences, Union City, CA, USA), quantified using a Quantus™ Fluorometer (Promega, USA) and libraries constructed. Sequencing was performed on the Illumina Miseq PE300 platform (Illumina, San Diego, USA) according to the standard protocol of Majorbio Bio-Pharm Technology Co. Ltd. (Shanghai, China).

### Sequencing data processing

Quality control of the double-ended raw sequencing data was performed using fastp v0.19.6 (Chen et al., 2018). Splicing was performed using FLASH v1.2.11 (Magoč and Salzberg, 2011): 1) truncate window bases for reads with an average quality score <20 within the 50 bp window of the tail, filter reads below 50 bp, and reads containing N bases were removed; 2) splice pairs of reads with a minimum overlap of 10 bp; 3) filter sequences with an overlap mismatch rate >0.2. Barcode and front-end primer sequences were removed. The data have been deposited in the Genome Sequence Archive (Tingting Chen et al., 2021) in National Genomics Data Center, China National Center (CNCB-NGDC Members and Partners, 2022) for Bioinformation / Beijing Institute of Genomics, Chinese Academy of Sciences (GSA: CRA010554) that are publicly accessible at https://ngdc.cncb.ac.cn/gsa.

Amplicon sequence variant (ASV) selection and feature table construction were performed using EasyAmplicon v1.14 (Liu et al., 2021): in the quality filter, a default value of 0.01 was used for “-fastq_maxee_rate”; in dereplication, a default value of 8 was used for “-miniuniqusize”; in denoising, “-minsize” was set to 20; in the feature table construction, the default statement was used; when removing plastids and non-bacteria, “-db” was set to rdp_16s_v18.fa, “-sintax_cutoff” was set to 0.6; normalization by subsample was set to the minimum value by default.

Alpha diversity was analyzed using EasyAmplicon: box plots and dilution curves were plotted using the default statements; filtering by abundance used a default value of 0 for “-thre”, true for “-scale” and 100 for “-zoom”; the default statement was used to filter the results above an abundance threshold (0.1%). The Venn network was plotted using EVenn (Tong Chen et al., 2021).

Beta diversity and relative abundance at the phylum and class level were analyzed using EasyAmplicon: default statements generated bray_curtis distance matrices, PCoA, CPCoA, and stackplot were plotted; in the treemap, “-topN” was set to 200; in the extended histogram, “-threshold” was set to the default value of 0.1, a default value of “t.test” was used for “--method”, a default value of 0.05 was used for the “--pvalue”, and “BH” was used for “--fdr”.

LEfSe analysis was performed using OECloud tools (https://cloud.oebiotech.cn.), with 0.1% abundance screening, unassigned categories were not included in the analysis, for the interclass Kruskal-Wallis test, alpha = 0.05, LDA score threshold was 2.0.

FAPROTAX v1.2.4 (Louca et al., 2016) in ImageGP (Chen et al., 2022) (http://www.ehbio.com/ImageGP/index.php/Home/Index/FAPROTAX.html) was used to predict biogeochemical cycle functional genes and STAMP v2.1.3 (Parks et al., 2014) was used to analyse for significant differences between the two groups: Welch’s t-test was used with a Storey FDR test correction. PICRUSt2 (Douglas et al., 2019) in the OECloud tools (https://cloud.oebiotech.cn) was used to predict the abundance of gene families. The volcano plot between the two groups was performed using EasyAmplicon and OmicStudio (Lyu et al., 2023) with “--threshold” set to 0, “--method” set to Wilcox, “--pvalue” set to the default value of 0.05 and “--fdr” set to 0.05. The top 25 enriched/depleted relative abundances in the volcano plot were plotted on a heatmap (R package ComplexHeatmap v2.15.1) (Gu et al., 2016; Gu, 2022).

The R package igraph v1.3.2 (Csardi and Nepusz, 2006) was used to analyze the network modules, calculate the Pearson correlation coefficient between two ASVs (occurrence >80%), maintain a threshold of r >0.7 and p <0.01 and calculate the layout based on layout_with_fr. The 18 modules with the highest number of nodes were colored.

Environmental factor analysis: ASVs (or KEGG orthologs) with station-wide frequencies ≥100 were screened for Hellinger transformation to attenuate the effect of zero values. Environmental factor data were +1 and then natural log transformed to make the data more homogeneous. Using the R package vegan v2.6-2 (Oksanen, 2010) decorana() operations for axis lengths and vif.cca() for covariance analysis, 999 permutation tests were carried out on the constraint axis as a whole and on each axis separately (Bonferroni for p-value correction). ordistep()/ordiR2step() was used for forward selection and 999 permutation tests, varpart() was used for variance decomposition followed by 999 permutation tests to explain variance. bioenv() was used to calculate the most relevant combination of environmental factors.

Random forest modelling was performed using the R package randomForest v4.7-1.1 with a default statement and with cross-validation to filter the number of features associated with temporal change (Liaw and Wiener, 2002; J. Zhang et al., 2018; Liu et al., 2021). The monthly affiliation of sediment samples was predicted based on the feature model, and a time-series prediction fit was plotted.

## Results

### Sequence screening and ASV characterization table

After quality control of 56 amplicon samples, 2,606,341 sequences were retained and 296,193 sequences were discarded. 6,723 good amplicons were obtained after redundancy and denoising. 6,497 ASVs were obtained after removal of plasmids and non-bacteria (1,754,033/2,606,341 = 67.3% sequences mapped). Sample size was 15,325 after equal sampling and normalization.

### Alpha diversity

Dilution curves by time series and by station and substrate type each reached a plateau (Fig. 1b). There were no significant differences in the Shannon indices between the monthly samples. The Shannon index for ZQ mud samples was significantly lower than that for ZQ sand samples (TukeyHSD, p = 0.001) and significantly lower than that for LC sand samples (p = 0.036) (Fig. 2b). There were 432 ASVs with a relative abundance >0.1% by station and substrate type, with 152-181 ASVs in each group. There were 284 ASVs that were unique to each group. There were 1,652 ASVs with abundances >0.1% when counted by temporal variation, with the least variation between samples collected from 5-7^th^ (December, February and March) and the most variation within samples collected from 8-10^th^ (April, May and July).

**Fig. 2.**
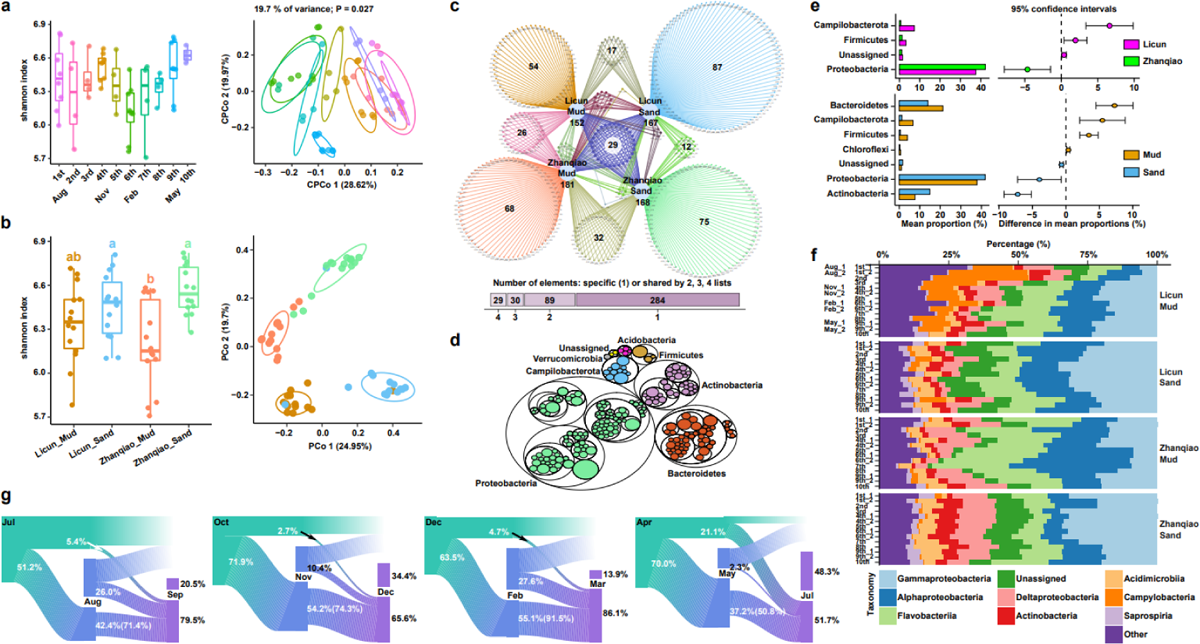
Alpha/Beta diversity and relative abundance at major phylum and class levels. a) Shannon index (TukeyHSD test) and CPCoA of samples by month b) Shannon index (TukeyHSD test) and PCoA (Bray_Curtis) of samples from different stations and substrates c) Venn network plots of bacterial ASVs (abundance >0.1%) between stations and substrate types d) Treemap of relative abundance top200 ASVs e) Comparison of differences in phylum levels (abundance >0.1%) between stations and between substrates (>0.5%, t.test, p <0.05) f) Relative abundance at major class levels g) Sankey plots of changes in bacterial ASVs (>0.1%) by season, with transmission rates of common ASVs in brackets.

### Beta diversity and relative abundance at major phylum and class levels

CPCoA analysis of samples from each month was significant at 19.7% resolution (p <0.05) and was best separated on CPCo axis 1. PCoA analyses separated samples from different stations and substrates (p <0.05). Most of the top200 ASVs by relative abundance belonged to Proteobacteria, Bacteroidetes and Actinobacteria. The relative abundance of Campilobacterota and Firmicutes was significantly higher at LC, while Proteobacteria was significantly higher at ZQ. Bacteroidetes, Campilobacterota, Firmicutes and Chloroflexi were significantly more abundant in the muddy samples, while Actinobacteria and Proteobacteria were significantly more abundant in the sandy samples. The composition at the class level varied between stations and substrates, although Gammaproteobacteria, Alphaproteobacteria, Flavobacteriia were most abundant with the sum of their relative abundances accounting for about 50% (Fig. 2).

### Linear Discriminant Analysis Effect Size (LEfSe) analysis

In the LEfSe of the monthly samples, Verrucomicrobia was the biomarker for the February sample, Bacilli and Verrucomicrobiae for February, Alphaproteobacteria for March, Bacteroidia for August and Cytophagia for October. Among the different stations and substrates, Acidobacteria and Bacteroidetes were biomarkers for the ZQ mud samples, Actinobacteria for ZQ sand, Campilobacterota and Firmicutes for LC mud, and Proteobacteria for LC sand (Fig. 3).

**Fig. 3.**
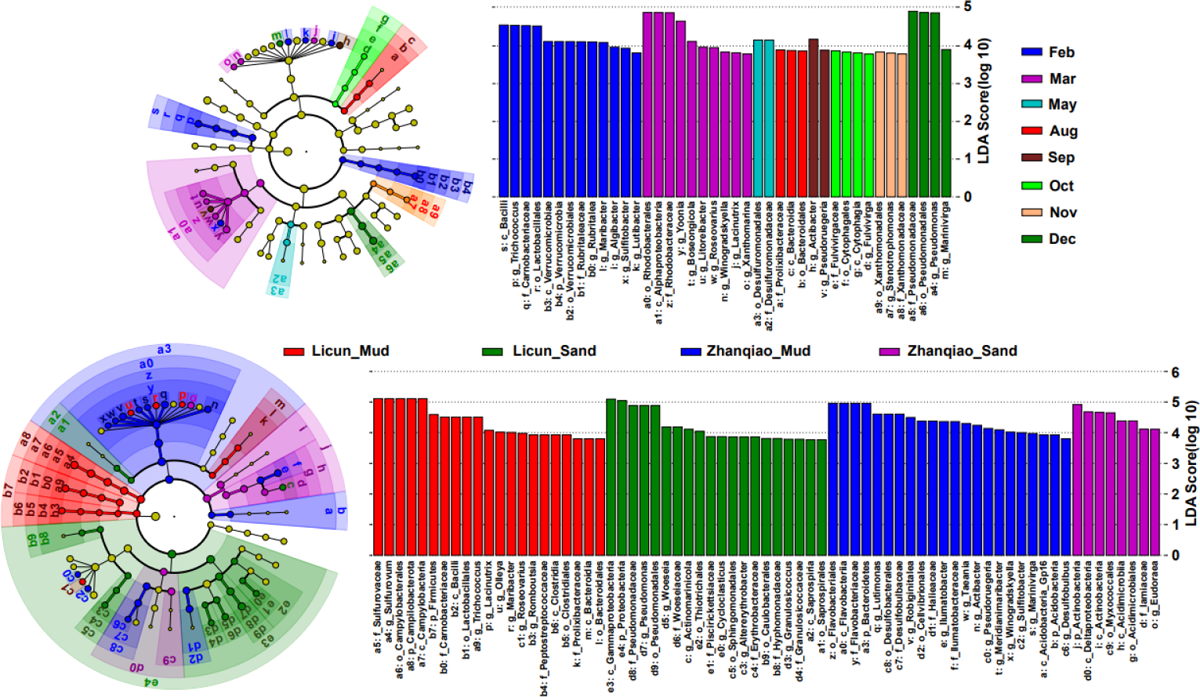
Time series and station/substrate LEfSe analysis. Dot diameters represent relative abundance (>0.1%, unassigned categories are not included in the analysis), no significant differences are yellow, Kruskal-Wallis tests for significant differences between classes (p <0.05) are assigned colors, LDA score threshold 2.0.

### Comparison of gene function

Chemoheterotrophy function was high at both stations (>30% in all), with significantly higher abundance at LC (34.6%, Welch’s t-test, p = 0.024). Respiration of sulfur compounds (11.4%) and sulfate respiration (11.2%) were also significantly higher at ZQ (p <0.001). The relative abundance of aerobic chemoheterotrophy was different between substrates, with sandy samples being highly significantly higher (29.7%, p = 0.0002). Fermentation abundance was significantly higher in mud samples (9.9%, p <0.001) (Fig. 4).

**Fig. 4.**
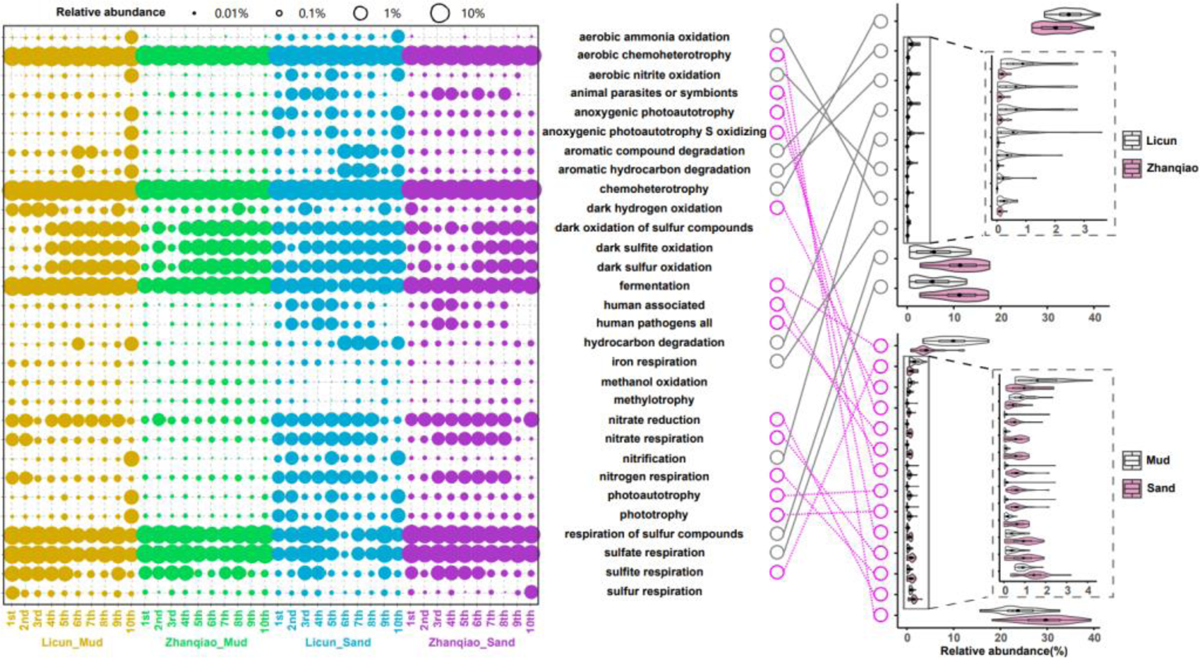
Relative abundance of elemental cycling functions (abundance >0.1%) and comparison of differences between stations and between substrates (abundance > 0.2%, Welch’s t-test, p < 0.05).

Among gene families, the relative abundance of those associated with lipid metabolism, terpenoid metabolism and polyketides was >0.5% and significantly higher at ZQ than at LC.

Between substrates, there were more significantly different KEGG_L2 (Kyoto Encyclopedia of Genes and Genomes, 16 categories with abundance >0.5%) involving metabolism, environmental information processing, cellular processes, organismal systems and human diseases. Most KEGG_L3 (90.2%) were not significantly different between stations, whereas 56.1% were significantly different between substrates (Fig. S3). KEGG orthologs (KO) was also mostly (81.7%) not significantly different between stations, although 64.5% were significantly different between substrates (Fig. 5).

**Fig. 5.**
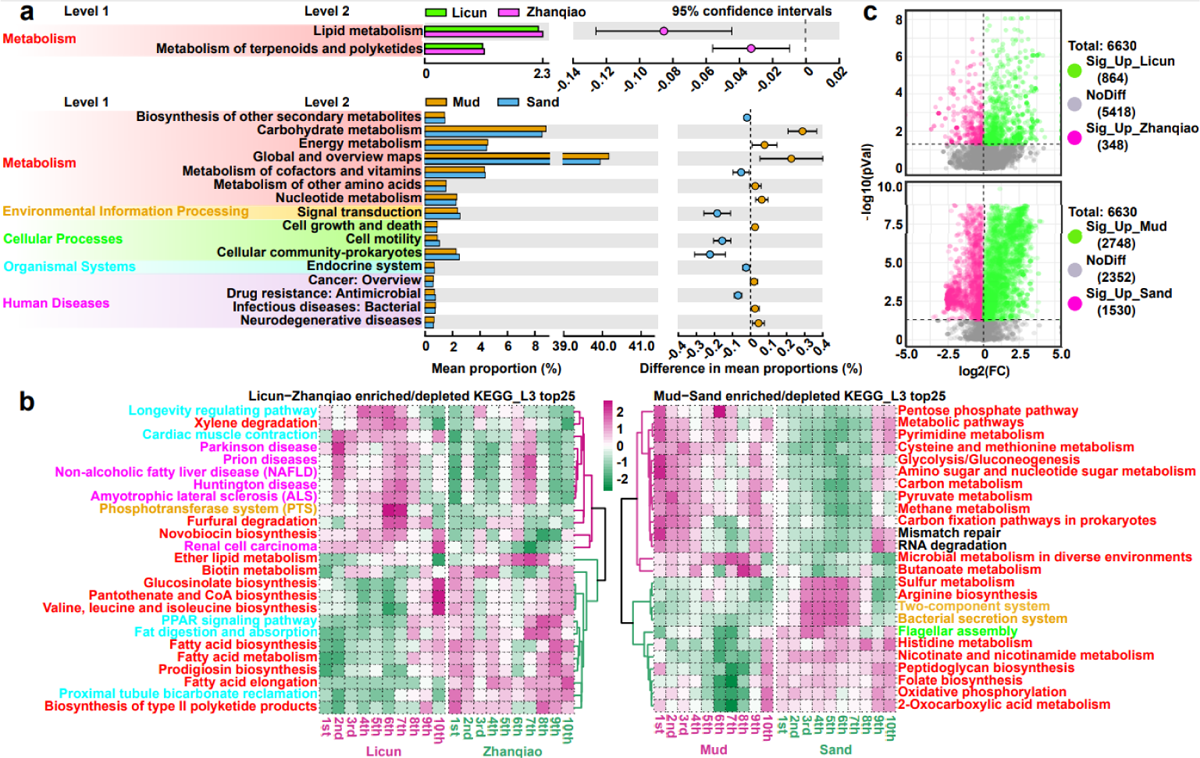
Gene family prediction. a) Comparison of KEGG differences between stations and substrates (abundance >0.5%, Welch’s t.test, adj <0.05) b) Relative abundance of major enriched/depleted KEGG_L3 between stations and substrates (top25, category names colored as in Fig. 5a KEGG_L1) c) KO volcanogram analysis across stations and substrates.

### Comparison of network modularity

The average edge number of nodes with significant correlations in summer, autumn, winter and spring were 12.3, 11.8, 16.6 and 21.9. Within stations, sandy samples have a higher number of nodes and a higher average connectivity degree than muddy samples (LC - muddy 8.6, sandy 11.4, ZQ - muddy 7.2, sandy 14.2) (Fig. 6).

**Fig. 6.**
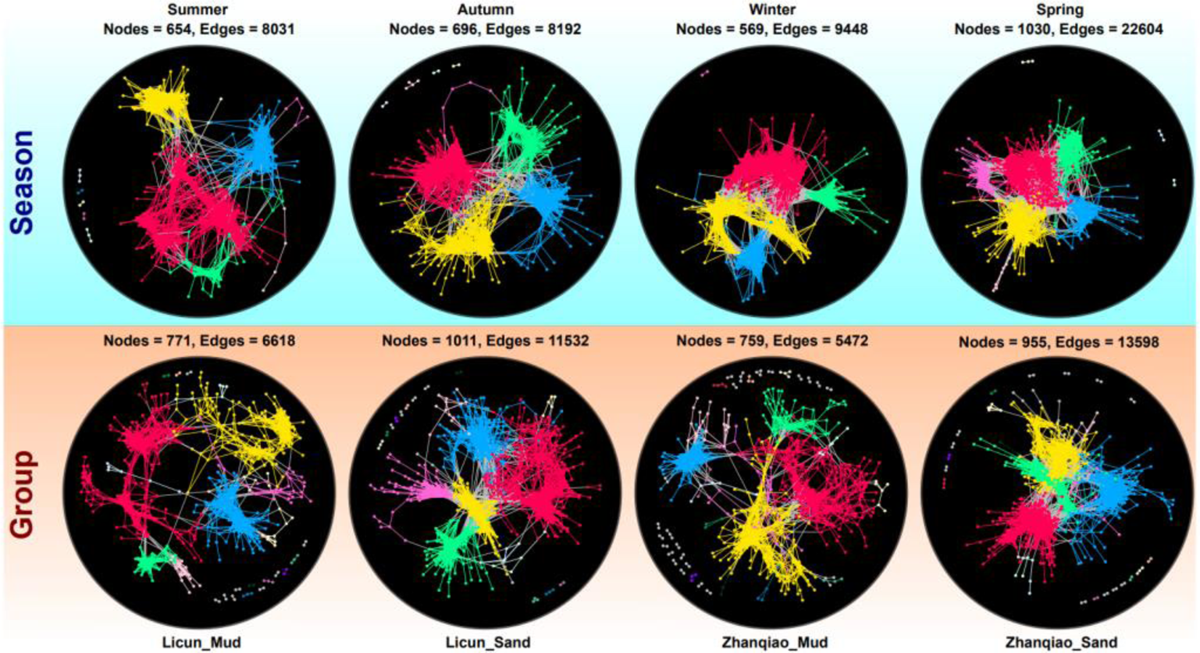
Network module analysis. The top row shows the network modules of different seasons; the bottom row shows the network modules of different stations and substrates. The Pearson correlation coefficient (r >0.7 and p <0.01) was calculated between two ASVs with an occurrence rate >80%. The layout was calculated based on layout_with_fr. In each network diagram, the top 18 modules are given a different color.

### Analysis of environmental factors

PIC + pH + DO% + PAR + Sal + Tem + POC + Chl were forward selected in this order. The total variance explained by PIC/pH/PAR was significant (p <0.05). The most relevant combination of environmental factors was PIC for size 1 (correlation 0.3154). On the RDA1 axis, there was a tendency for samples to be separated by season (Fig. 7).

**Fig. 7.**
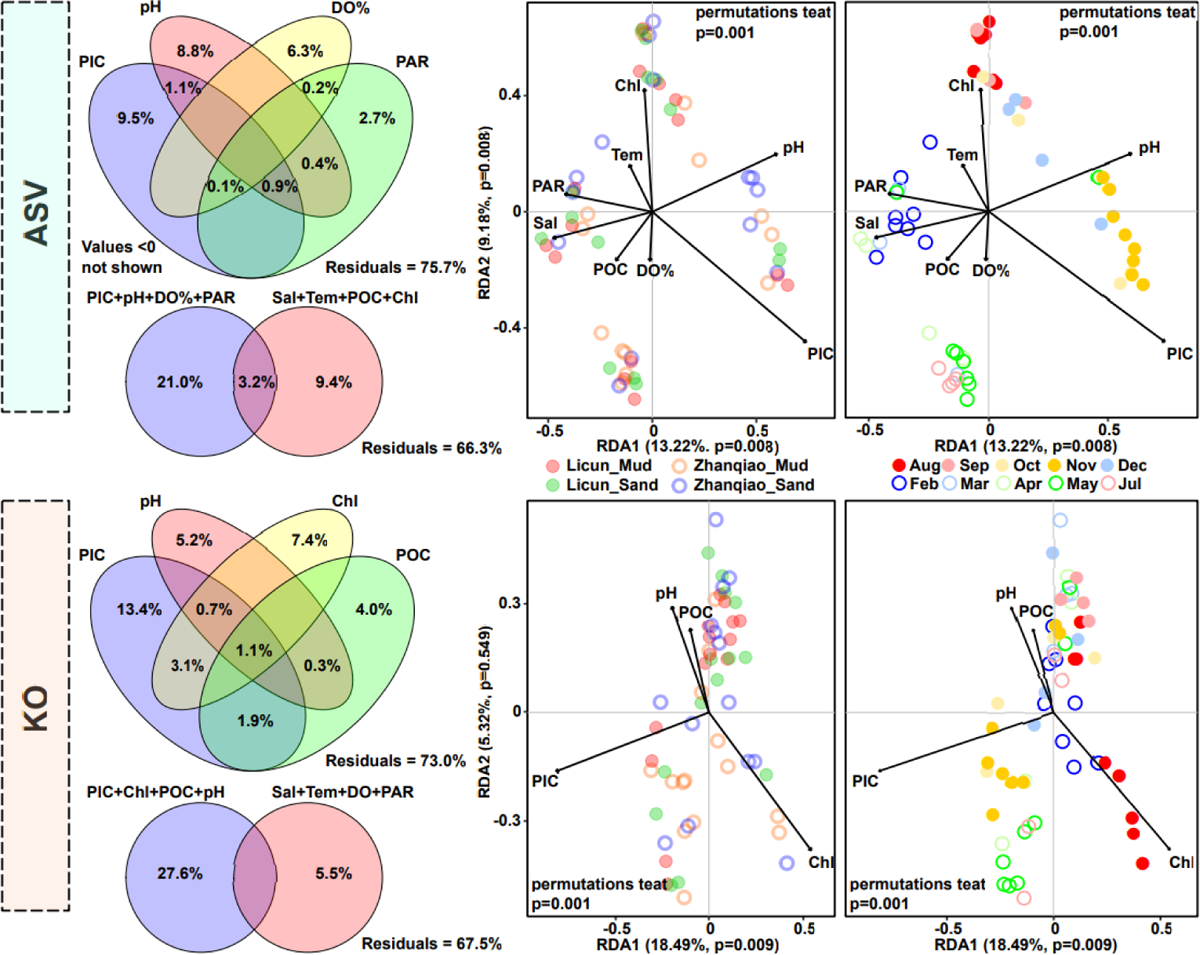
Environmental factor analysis. Individual and combined variance decomposition of the best four factors in forward selection, parsimonious model after forward selection showing the first two axes of the RDA (type II scale, double ordinate plot, sample coordinates calculated using species weights). Tem: water temperature DO%: dissolved oxygen air saturation Sal: salinity PAR: photosynthetically active radiation PIC: particulate inorganic carbon POC: particulate organic carbon.

PIC + pH + Chl + POC passed forward selection in order and the total variance explained by PIC/Chl was significant (p <0.01). The most relevant environmental factor combination was Chl + PIC for size 2 (correlation 0.3103), followed by Chl + PIC + POC for size 3 (0.2870) and Chl for size 1 (0.2694). There was less variation between samples compared to the results of the ASV redundancy analysis (Fig. 7).

### Random forest prediction

Cross-validation yielded 43 family-level features that were the better combinations, with the Proteobacteria of spongiibacteraceae, porticoccaceae, granulosicoccaceae, alcaligenaceae, nannocystaceae and methylophilaceae having the highest importance (%IncMSE) (Fig. S5).

## Discussion

### Relationship between intertidal sediment biofilms by station and substrate factors

The Shannon index did not differ significantly between stations, but was less variable and significantly higher in sandy samples than in muddy samples (Fig. S1). A comparison of network modularity also showed that the correlations between the ASVs were tighter and that there were more ASVs involved in community assembly in sandy samples than in mud (Fig. 6). Network complexity and connectivity are often positively correlated with environmental heterogeneity and microbial communities in heterogeneous environments are more likely to aggregate and form cooperative networks to share resources and transfer information (Mougi and Kondoh, 2012; Morriën et al., 2017; Lu et al., 2022). It is hypothesized here that differences in current strength and runoff within and outside the bay may have had a stronger effect on permeable sandy sediments, with increased heterogeneity promoting bacterial aggregation to adapt to the environment.

Of the highly abundant bacteria in each station/substrate type, ASVs unique to each group accounted for 35.5-52.1% of the groups, indicating variability in the composition of high abundance bacteria in each sample group, even between samples of different substrates sampled in close proximity within stations. Bacterial diversity has been found to differ between fine- and coarse-grained sediments (Lai et al., 2023). Differences in sediment grain structure can also lead to changes in bacterial community structure (van de Kamp et al., 2019). The abundance of nitrification-associated bacteria fluctuates more in intertidal sandy sediments than in mud (Fernandes et al., 2016). Sediment grain size has a significant effect on the community structure of *nirS*-type denitrifying bacteria (Chen et al., 2020). Therefore, sediments of different grain sizes have the ability to shape their own specific bacterial communities.

Proteobacteria were the most abundant group in this study, accounting for about 50% (Fig. S2). They were slightly less abundant in LC mud samples (especially in summer, when Campilobacterota appeared in high abundance, increasing from an average of 1.2% to 14.6-34.1%). Proteobacteria are widely distributed, have high metabolic, cooperative and biofilm-forming capacities, and are also adapted to high metal loads (Regnell and Watras, 2019; Zhuang et al., 2019; Hillyer et al., 2023). Proteobacteria have been detected in high abundance in estuarine intertidal sediments (37.5%) (Yi et al., 2020), coral reef sediments (65%) (Alvarez-Yela et al., 2019), mangrove surface sediments (>40%) (Huergo et al., 2018; Fiard et al., 2022), polar surface sediments (44-68%) (Thomas et al., 2020; Chaudhary et al., 2022), and marine sediments exposed to oil transport activities (45-77%) (Oyetibo et al., 2021). Campilobacterota has been observed at higher abundances (>5%) in thermogenic hydrocarbon seep sediments (Li et al., 2023) and is involved in mediating oxidation of sulfur, sulfide or sulphate in intertidal sediments (7.90-22.15%) (Carrier et al., 2020; Fang et al., 2022). The genus Arcobacter has been found to increase in abundance in anoxic surface sediments, and in addition, is highly responsive to oxygenation events (Broman et al., 2017; Mori et al., 2021). Therefore, it is likely that hydrocarbon, sulfide and oxygen (low summer oxygen, low mud permeability and low circulation rates within the bay) are responsible for the increased abundance of Campilobacterota in summer mud samples at LC.

There are corresponding phylum-level biomarkers in different stations and substrates. All taxonomic levels of Firmicutes and Campilobacterotas were biomarkers in LC mud samples, Firmicutes is one of the major groups in natural biofilms (Guo et al., 2017; Liang et al., 2019) and is abundant in polyethylene microplastic biofilms, and may provide potential hosts for antibiotic resistance genes (Wu et al., 2017; Guo et al., 2020). Firmicutes is also highly abundant in environments under severe stress, such as in deep-sea sediments (9.5%) (Goffredi and Orphan, 2010; Franco et al., 2020), shallow hypoxic zone sediments (Bhattacharya et al., 2021), and anaerobic bioreactors (involved in hydrolysis and fermentation) (Narihiro et al., 2015; Fontana et al., 2016; Luo et al., 2016). The abundance of all taxonomic levels of Acidobacteria increased in ZQ mud samples and most were also increased in ZQ sand samples. Acidobacteria are low-nutrient bacteria (Fierer et al., 2007) that have been found to be adapted to polar environments (Pearce et al., 2013; Gugliandolo et al., 2016) and are key microorganisms (∼8%) in mangrove sediments (Huergo et al., 2018; Tavares et al., 2021; Fiard et al., 2022). Actinobacteria is one of the major bacterial groups in estuarine surface sediments and is significantly and positively correlated with denitrification rates (Wei et al., 2015; Li et al., 2022). Uncultured Actinobacteria OPB41 has been shown to be adapted to deep-sea sediment environments, where it can catabolize sugars and be metabolically active (Zinke et al., 2017; Bird et al., 2019). In summary, there were significant differences in the abundance patterns of the more diverse bacterial taxa and functions between the different stations and substrates, shaping their own specific bacterial communities.

Chemoheterotrophy function occurred in 77.8% (900 of 1,157 ASVs) and aerobic chemoheterotrophy function occurred in 58.5%. One study has found that the abundance of chemoheterotrophy and aerobic chemoheterotrophy in surface sediments may account for up to half of the total abundance, giving bacteria a strong ability to degrade organic matter and obtain energy from the oxidation of organic compounds (X. Zhang et al., 2018; Hou et al., 2021). Chemoheterotrophy and aerobic chemoheterotrophy account for more than 50% of the bacteria and probably point to a stronger tendency for organic matter degradation in LC or sandy sediments. Sulfur compounds respiration function occurred in 26.4% (306 of 1,157 ASVs) and sulfate respiration function occurred in 24.4% (282 of 1,157 ASVs). Sulfur compounds respiration may be associated with adaptation to hypoxia by promoting respiration using sulfides as electron acceptors (Deng et al., 2019). Sulfate respiration may play a major role in the biodegradation of sulfate-dependent aromatic compounds (Zhuang et al., 2019). Fermentation function occurred in 22.0% (255 of 1,157 ASVs). It has been found that microorganisms in intertidal and subtidal sandy sediments often switch between oxic and anoxic states, favouring facultative anaerobic fermenters where fermentation is dominated by anoxic carbon mineralization (which also promotes H2 production), in contrast to muddy sediments where it is largely unrelated to anaerobic respiration (Precht et al., 2004; Kessler et al., 2013; Saad et al., 2017; Kessler et al., 2019).

Fermentation also promotes the cycling of other material, such as in organic matter-rich and sulfate-rich environments, where enhanced fermentation may lead to increased microbial reduction of sulfate (Sun et al., 2020). The abundant biogeochemical cycling functions differ significantly between stations and substrates, with stations mostly differing significantly in chemoheterotrophy, sulphur, nitrogen, iron, aromatics and hydrocarbon metabolism, whereas substrates differ in aerobic chemoheterotrophy, fermentation, dark hydrogen oxidation, animal-related, photosynthesis-related and sulphite metabolism, suggesting that both station and substrate factors are likely to be differentially involved in biogeochemical cycling processes between samples.

There were significantly fewer categories of significant differences between stations than between substrates in KEGG_L2, suggesting that differences in gene functions were more due to substrate differences than to differences in location within and outside the bay. Between stations, the enriched/depleted KEGG_L3 top25 involved more organismal systems and human diseases (Fig. 5b), with all human disease categories enriched at LC, suggesting that the poor circulation in the bay was not conducive to effective dispersal of land-based sources of domestic sewage, etc. Between the substrates, the top 25 gene functions were mostly related to metabolism. Significant differences in KO PCoA were found between LC substrates and between muddy samples (Fig. S4). ZQ current velocities subjected the sediments to strong scouring, which promoted similar bacterial gene family succession for both muddy and sandy samples. The low current velocities in the bay may have been sufficient to influence bacterial succession in the more permeable sandy sediments, so that there were no gene family differences between stations for the sandy samples. In contrast, the sediment environment had less influence on gene families than on ASVs, possibly indicating that the bacterial community was evolving but the core gene functions adapted to the environment were relatively stable.

### Temporal changes in intertidal sediment biofilms

Temporal changes in bacterial community structure varies between intertidal and supratidal environments in different areas, with some showing no significant changes and others showing significant shifts (Taylor and Kurtz, 2020). Seasonal changes in bacterial communities are sometimes not as large as spatial changes, when seasonal changes are only apparent in individual taxa (Dini-Andreote et al., 2014; Tebbe et al., 2022). In contrast, bacterioplankton in the Chesapeake Bay have a repeating annual pattern, with strong temporal variation in bacterial communities even overwhelming spatial patterns (Wang et al., 2020). In other estuarine and coastal waters, temporal variation also strongly influences changes in bacterial abundance (Lindh et al., 2015; Bunse and Pinhassi, 2017; Thomson et al., 2022). Bacterioplankton in estuarine wetlands have a high diversity index in summer, and DO and pH strongly influence these communities (Li et al., 2020). In another study of mangrove and tidal wetland sediments, bacterial diversity indices were also higher in summer (Zhou et al., 2017). Dry-rainy season variation was the dominant factor influencing these communities, with temporal and spatial patterns more pronounced in water column bacterial communities than in sediments (Kaestli et al., 2017). In another study, intertidal sediment bacteria were significantly separated in PCoA when grouped by wet and dry seasons (Yi et al., 2020). There is also indirect evidence for the influence of temporal variation on bacterial communities: acyl-homoserine lactones (AHLs), which are key bacterial signals within biofilms that are degraded by pH- and temperature-dependent hydrolysis, vary seasonally in concentration, being higher in winter and lower in summer (Roggatz and Parsons, 2022). The intertidal filter-feeding clam (*Ruditapes philippinarum*) is closely related to the microorganisms in its environment, and its extrapallial fluids show strong temporal variation in bacterial alpha-diversity between months (Offret et al., 2023).

Of the most abundant bacteria, the least variation over time was seen in winter and the greatest variation in spring (Fig. 2g). Samples collected from April to July were also the most distinct from each other in the CPCoA. In the network modularity comparison, winter and spring samples were more closely correlated than summer and autumn ASVs, with the lowest number of nodes in winter (569) and the highest in spring (1,030) (Fig. 6). Based on the projection of samples on environmental factors by month in the RDA, it is clear that spring samples (April to July samples) were most positively influenced by POC, DO%, PIC and Sal (and most negatively influenced by Tem/Chl/PH), summer samples (August and September samples) were positively influenced by Chl/Tem (and negatively influenced by PIC/DO%/POC), autumn samples (October to December samples) were positively influenced by pH/PIC (and negatively influenced by Sal/PAR) and winter samples (February and March samples) were positively influenced by PAR/Sal (and negatively influenced by pH) (Fig. 7). The spring season was affected by a combination of more environmental factors and the abundant bacteria correlated with each environmental factor varied. The network diagram suggests that the most species were involved in community building in spring, so there was a tendency for greater species variation with environmental changes, whereas in winter the bacteria were more closely related, but had the least number of community building species, so the tendency for variation might be minimal.

Monthly samples generally had their own biomarkers at the order level: August - Bacteroidales, October - Cytophagales, November - Xanthomonadales, December - Pseudomonadales, February - Verrucomicrobiales and Lactobacillales, March - Rhodobacterales, and May - Desulfuromonadales. It can be seen that the temporal change had a strong influence in shaping the intertidal bacterial community in the temperate Jiaozhou Bay, and there was a tendency for taxonomic abundance at the order level to change from month to month. The pattern of change in abundance of the better combination in the random forest prediction responded better to changes in temporal change (rho > 0.7, p < 0.001, R^2^ > 0.6), reflecting the effect of temporal change on bacterial communities from the other side.

The trend of seasonal separation of ASV in the RDA analysis was reduced in the KO analysis. Similar results were also found for CPCoA (Fig. S4). It has previously been reported that when there were seasonal changes in the bacterial community structure in the Monterey Bay upwelling zone, there was a consistent seasonal trend in the gene abundance of key metabolic pathways (Reji et al., 2020). Bacterial communities in surface waters of Kuwait Bay and nearby coastal waters showed significant seasonal separation, while PICRUSt functional predictions showed partial seasonality (Ismail and Almutairi, 2022). Thus, temporal change had a major influence in the Qingdao intertidal bacterial community, but seems to have had less influence in shaping the gene family of the community.

## Conclusion

The analyses presented here show that sandy intertidal sediments have more bacterial species involved in community building than muddy sediments. Greater current strengths over the permeable sandy sediments and high environmental stress increased heterogeneity, which in turn promoted bacterial community aggregation and their ability to adapt to the environment. There was a significant separation between substrate types in the PCo 1 axis and between stations in the PCo 2 axis. Different substrate types at the different stations had their own phylum-level biomarkers and bacterial functional enrichment status, with substrate factors having a greater influence. Notably, many gene families associated with human disease were enriched at LC, suggesting that the low circulation rates in the bay were unable to remove contamination from the sources of domestic wastewater. The sediment environment had less influence on KO than on ASV, and it is possible that bacterial species are constantly evolving, but the function of genes adapted to the environment is relatively stable. Temporal changes had a stronger shaping effect on the composition of bacterial species than on gene families. These results will provide the necessary empirical support for ecological conservation in temperate intertidal sediments.

## Author contributions

**Xuechao Chen, Xinran Zhang, Hao Yu**: Methodology, Software, Validation, Formal analysis, Investigation, Project administration, Visualization, Writing - original draft. **Meiaoxue Han, Jianhua Sun, Gang Liu**: Software, Investigation, Writing - review and editing. **Yan Ji, Chuan Zhai, Liyan Zhu**: Writing - review and editing. **Hongbing Shao**: Resources, Supervision. **Min Wang, Andrew McMinn, Yantao Liang**: Conceptualization, Resources, Funding acquisition, Supervision, Writing - review and editing.

## Acknowledgements

We are grateful for the use of the high-performance server at the Center for High Performance Computing and System Simulation, Pilot National Laboratory for Marine Science and Technology (Qingdao, China). We appreciate the computational resources provided by IEMB-1, a high-performance computing cluster operated by the Institute of Evolution and Marine Biodiversity, and the Marine Big Data Center of the Institute for Advanced Ocean Study of Ocean University of China. The data were also analyzed on the free online platform of Majorbio I-Sanger Cloud Platform (www.i-sanger.com) and OECloud tools (https://cloud.oebiotech.cn).

This work was supported by the Laoshan Laboratory (No. LSKJ202203201), National Key Research and Development Program of China (2022YFC2807500), Natural Science Foundation of China (No. 42120104006, 41976117 and 42176111) and the Fundamental Research Funds for the Central Universities (202172002, 201812002, 202072001 and Andrew McMinn).

## References

Alvarez-Yela, A.C., Mosquera-Rendón, J., Noreña-P, A., Cristancho, M., López-Alvarez, D., 2019. Microbial diversity exploration of marine hosts at Serrana Bank, a coral atoll of the Seaflower Biosphere Reserve. Front. Mar. Sci. 6, 338. https://doi.org/10.3389/fmars.2019.00338

Avila-Jimenez, M.-L., Burns, G., He, Z., Zhou, J., Hodson, A., Avila-Jimenez, J.-L., Pearce, D., 2020. Functional associations and resilience in microbial communities. Microorganisms 8, 951. https://doi.org/10.3390/microorganisms8060951

Bhattacharya, S., Mapder, T., Fernandes, S., Roy, C., Sarkar, J., Rameez, M.J., Mandal, S., Sar, A., Chakraborty, A.K., Mondal, N., Chatterjee, S., Dam, B., Peketi, A., Chakraborty, R., Mazumdar, A., Ghosh, W., 2021. Sedimentation rate and organic matter dynamics shape microbiomes across a continental margin. Biogeosciences 18, 5203–5222. https://doi.org/10.5194/bg-18-5203-2021

Bird, J.T., Tague, E.D., Zinke, L., Schmidt, J.M., Steen, A.D., Reese, B., Marshall, I.P.G., Webster, G., Weightman, A., Castro, H.F., Campagna, S.R., Lloyd, K.G., 2019. Uncultured microbial phyla suggest mechanisms for multi-thousand-year subsistence in Baltic Sea sediments. mBio 10, e02376–18. https://doi.org/10.1128/mBio.02376-18

Broman, E., Sachpazidou, V., Pinhassi, J., Dopson, M., 2017. Oxygenation of hypoxic coastal Baltic Sea sediments impacts on chemistry, microbial community composition, and metabolism. Front. Microbiol. 8, 2453. https://doi.org/10.3389/fmicb.2017.02453

Bunse, C., Pinhassi, J., 2017. Marine bacterioplankton seasonal succession dynamics. Trends Microbiol. 25, 494–505. https://doi.org/10.1016/j.tim.2016.12.013

Carrier, V., Svenning, M.M., Gründger, F., Niemann, H., Dessandier, P.-A., Panieri, G., Kalenitchenko, D., 2020. The impact of methane on microbial communities at marine arctic gas hydrate bearing sediment. Front. Microbiol. 11, 1932. https://doi.org/10.3389/fmicb.2020.01932

Chaudhary, D.K., Karki, H.P., Bajagain, R., Kim, H., Rhee, T.S., Hong, J.K., Han, S., Choi, Y.-G., Hong, Y., 2022. Mercury and other trace elements distribution and profiling of microbial community in the surface sediments of East Siberian Sea. Mar. Pollut. Bull. 185, 114319. https://doi.org/10.1016/j.marpolbul.2022.114319

Chen, Q., Fan, J., Ming, H., Su, J., Wang, Y., Wang, B., 2020. Effects of environmental factors on denitrifying bacteria and functional genes in sediments of Bohai Sea, China. Mar. Pollut. Bull. 160, 111621. https://doi.org/10.1016/j.marpolbul.2020.111621

Chen, S., Zhou, Y., Chen, Y., Gu, J., 2018. fastp: an ultra-fast all-in-one FASTQ preprocessor. Bioinformatics 34, i884–i890. https://doi.org/10.1093/bioinformatics/bty560

Chen, Tingting, Chen, X., Zhang, S., Zhu, J., Tang, B., Wang, A., Dong, L., Zhang, Zhewen, Yu, C., Sun, Yanling, Chi, L., Chen, H., Zhai, S., Sun, Yubin, Lan, L., Zhang, X., Xiao, J., Bao, Y., Wang, Y., Zhang, Zhang, Zhao, W., 2021. The Genome Sequence Archive Family: Toward Explosive Data Growth and Diverse Data Types. Genomics Proteomics Bioinformatics 19, 578–583. https://doi.org/10.1016/j.gpb.2021.08.001

Chen, T., Liu, Y., Huang, L., 2022. ImageGP: An easy-to-use data visualization web server for scientific researchers. iMeta 1. https://doi.org/10.1002/imt2.5

Chen, Tong, Zhang, H., Liu, Y., Liu, Y.-X., Huang, L., 2021. EVenn: Easy to create repeatable and editable Venn diagrams and Venn networks online. J. Genet. Genomics 48, 863–866. https://doi.org/10.1016/j.jgg.2021.07.007

CNCB-NGDC Members and Partners, 2022. Database resources of the national genomics data center, china national center for bioinformation in 2022. Nucleic Acids Res. 50, D27–D38. https://doi.org/10.1093/nar/gkab951

Csardi, G., Nepusz, T., 2006. The igraph software package for complex network research. InterJournal Complex Syst. 1695, 1–9.

Dai, J., Song, J., Li, X., Yuan, H., Li, N., Zheng, G., 2007. Environmental changes reflected by sedimentary geochemistry in recent hundred years of Jiaozhou Bay, North China. Environ. Pollut. 145, 656–667. https://doi.org/10.1016/j.envpol.2006.10.005

Dang, H., Lovell, C.R., 2016. Microbial surface colonization and biofilm development in marine environments. Microbiol. Mol. Biol. Rev. 80, 91–138. https://doi.org/10.1128/MMBR.00037-15

Deng, J., Auchtung, J.M., Konstantinidis, K.T., Brettar, I., Höfle, M.G., Tiedje, J.M., 2019. Genomic variations underlying speciation and niche specialization of *Shewanella baltica*. mSystems 4, e00560–19. https://doi.org/10.1128/mSystems.00560-19

Dini-Andreote, F., de Cássia Pereira e Silva, M., Triadó-Margarit, X., Casamayor, E.O., van Elsas, J.D., Salles, J.F., 2014. Dynamics of bacterial community succession in a salt marsh chronosequence: evidences for temporal niche partitioning. ISME J. 8, 1989–2001. https://doi.org/10.1038/ismej.2014.54

Douglas, G.M., Maffei, V.J., Zaneveld, J., Yurgel, S.N., Brown, J.R., Taylor, C.M., Huttenhower, C., Langille, M.G.I., 2019. PICRUSt2: An improved and customizable approach for metagenome inference (preprint). Bioinformatics. https://doi.org/10.1101/672295

Duarte, B., Freitas, J., Caçador, I., 2012. Sediment microbial activities and physic-chemistry as progress indicators of salt marsh restoration processes. Ecol. Indic. 19, 231–239. https://doi.org/10.1016/j.ecolind.2011.07.014

Fang, J., Jiang, W., Meng, S., He, W., Wang, G., Guo, E., Yan, Y., 2022. Polychaete bioturbation alters the taxonomic structure, co-occurrence network, and functional groups of bacterial communities in the intertidal flat. Microb. Ecol. https://doi.org/10.1007/s00248-022-02036-2

Fernandes, S.O., Javanaud, C., Michotey, V.D., Guasco, S., Anschutz, P., Bonin, P., 2016. Coupling of bacterial nitrification with denitrification and anammox supports N removal in intertidal sediments (Arcachon Bay, France). Estuar. Coast. Shelf Sci. 179, 39–50. https://doi.org/10.1016/j.ecss.2015.10.009

Fiard, M., Cuny, P., Sylvi, L., Hubas, C., Jézéquel, R., Lamy, D., Walcker, R., El Houssainy, A., Heimbürger-Boavida, L.-E., Robinet, T., Bihannic, I., Gilbert, F., Michaud, E., Dirberg, G., Militon, C., 2022. Mangrove microbiota along the urban-to-rural gradient of the Cayenne estuary (French Guiana, South America): Drivers and potential bioindicators. Sci. Total Environ. 807, 150667. https://doi.org/10.1016/j.scitotenv.2021.150667

Fierer, N., Bradford, M.A., Jackson, R.B., 2007. Toward an ecological classification of soil bacteria. Ecology 88, 1354–1364. https://doi.org/10.1890/05-1839

Fontana, A., Patrone, V., Puglisi, E., Morelli, L., Bassi, D., Garuti, M., Rossi, L., Cappa, F., 2016. Effects of geographic area, feedstock, temperature, and operating time on microbial communities of six full-scale biogas plants. Bioresour. Technol. 218, 980–990. https://doi.org/10.1016/j.biortech.2016.07.058

Franco, N.R., Giraldo, M.Á., López-Alvarez, D., Gallo-Franco, J.J., Dueñas, L.F., Puentes, V., Castillo, A., 2020. Bacterial composition and diversity in deep-sea sediments from the Southern Colombian Caribbean Sea. Diversity 13, 10. https://doi.org/10.3390/d13010010

Ge, Y., Lou, Y., Xu, M., Wu, C., Meng, J., Shi, L., Xia, F., Xu, Y., 2021. Spatial distribution and influencing factors on the variation of bacterial communities in an urban river sediment. Environ. Pollut. 272, 115984. https://doi.org/10.1016/j.envpol.2020.115984

Goffredi, S.K., Orphan, V.J., 2010. Bacterial community shifts in taxa and diversity in response to localized organic loading in the deep sea. Environ. Microbiol. 12, 344–363. https://doi.org/10.1111/j.1462-2920.2009.02072.x

Grabowski, R.C., Droppo, I.G., Wharton, G., 2011. Erodibility of cohesive sediment: The importance of sediment properties. Earth-Sci. Rev. 105, 101–120. https://doi.org/10.1016/j.earscirev.2011.01.008

Gu, Z., 2022. Complex heatmap visualization. iMeta 1. https://doi.org/10.1002/imt2.43

Gu, Z., Eils, R., Schlesner, M., 2016. Complex heatmaps reveal patterns and correlations in multidimensional genomic data. Bioinformatics 32, 2847–2849. https://doi.org/10.1093/bioinformatics/btw313

Gugliandolo, C., Michaud, L., Lo Giudice, A., Lentini, V., Rochera, C., Camacho, A., Maugeri, T.L., 2016. Prokaryotic community in lacustrine sediments of byers peninsula (Livingston Island, Maritime Antarctica). Microb. Ecol. 71, 387–400. https://doi.org/10.1007/s00248-015-0666-8

Guo, X., Niu, Z., Lu, D., Feng, J., Chen, Y., Tou, F., Liu, M., Yang, Y., 2017. Bacterial community structure in the intertidal biofilm along the Yangtze Estuary, China. Mar. Pollut. Bull. 124, 314–320. https://doi.org/10.1016/j.marpolbul.2017.07.051

Guo, X., Sun, X., Chen, Y., Hou, L., Liu, M., Yang, Y., 2020. Antibiotic resistance genes in biofilms on plastic wastes in an estuarine environment. Sci. Total Environ. 745, 140916. https://doi.org/10.1016/j.scitotenv.2020.140916

Hillyer, K.E., Raes, E., Bissett, A., Beale, D.J., 2023. Multi-omics eco-surveillance of bacterial community function in legacy contaminated estuary sediments. Environ. Pollut. 318, 120857. https://doi.org/10.1016/j.envpol.2022.120857

Hou, Y., Li, B., Feng, G., Zhang, C., He, J., Li, H., Zhu, J., 2021. Responses of bacterial communities and organic matter degradation in surface sediment to *Macrobrachium nipponense* bioturbation. Sci. Total Environ. 759, 143534. https://doi.org/10.1016/j.scitotenv.2020.143534

Huergo, L.F., Rissi, D.V., Elias, A.S., Gonçalves, M.V., Gernet, M.V., Barreto, F., Dahmer, G.W., Reis, R.A., Pedrosa, F.O., Souza, E.M., Monteiro, R.A., Baura, V.A., Balsanelli, E., Cruz, L.M., 2018. Influence of ancient anthropogenic activities on the mangrove soil microbiome. Sci. Total Environ. 645, 1–9. https://doi.org/10.1016/j.scitotenv.2018.07.094

Ismail, N., Almutairi, A., 2022. Bacterioplankton community profiling of the surface waters of Kuwait. Front. Mar. Sci. 9, 838101. https://doi.org/10.3389/fmars.2022.838101

Kaestli, M., Skillington, A., Kennedy, K., Majid, M., Williams, D., McGuinness, K., Munksgaard, N., Gibb, K., 2017. Spatial and temporal microbial patterns in a tropical macrotidal estuary subject to urbanization. Front. Microbiol. 8, 1313. https://doi.org/10.3389/fmicb.2017.01313

Kallmeyer, J., Pockalny, R., Adhikari, R.R., Smith, D.C., D’Hondt, S., 2012. Global distribution of microbial abundance and biomass in subseafloor sediment. Proc. Natl. Acad. Sci. 109, 16213–16216. https://doi.org/10.1073/pnas.1203849109

Kessler, A.J., Chen, Y.-J., Waite, D.W., Hutchinson, T., Koh, S., Popa, M.E., Beardall, J., Hugenholtz, P., Cook, P.L.M., Greening, C., 2019. Bacterial fermentation and respiration processes are uncoupled in anoxic permeable sediments. Nat. Microbiol. 4, 1014–1023. https://doi.org/10.1038/s41564-019-0391-z

Kessler, A.J., Glud, R.N., Cardenas, M.B., Cook, P.L.M., 2013. Transport zonation limits coupled nitrification-denitrification in permeable sediments. Environ. Sci. Technol. 47, 13404–13411. https://doi.org/10.1021/es403318x

Lai, X., Li, X., Song, J., Yuan, H., Duan, L., Li, N., Wangi, Y., 2023. Nitrogen loss from the coastal shelf of the East China Sea: Implications of the organic matter. Sci. Total Environ. 854, 158805. https://doi.org/10.1016/j.scitotenv.2022.158805

Li, C., Adebayo, O., Ferguson, D.K., Wang, S., Rattray, J.E., Fowler, M., Webb, J., Campbell, C., Morrison, N., MacDonald, A., Hubert, C.R.J., 2023. Bacterial anomalies associated with deep sea hydrocarbon seepage along the Scotian Slope. Deep Sea Res. Part Oceanogr. Res. Pap. 193, 103955. https://doi.org/10.1016/j.dsr.2022.103955

Li, M., Mi, T., Yu, Z., Ma, M., Zhen, Y., 2020. Planktonic bacterial and archaeal communities in an artificially irrigated estuarine wetland: diversity, distribution, and responses to environmental parameters. Microorganisms 8, 198. https://doi.org/10.3390/microorganisms8020198

Li, M., Wei, G., Liu, J., Wang, X., Hou, L., Gao, Z., 2022. Effects of nitrate exposure on nitrate reduction processes in the wetland sediments from the Yellow River estuary. Estuaries Coasts 45, 315–330. https://doi.org/10.1007/s12237-021-00966-7

Liang, X., Peng, L.-H., Zhang, S., Zhou, S., Yoshida, A., Osatomi, K., Bellou, N., Guo, X.-P., Dobretsov, S., Yang, J.-L., 2019. Polyurethane, epoxy resin and polydimethylsiloxane altered biofilm formation and mussel settlement. Chemosphere 218, 599–608. https://doi.org/10.1016/j.chemosphere.2018.11.120

Liaw, A., Wiener, M., 2002. Classification and regression by randomForest. R News 2, 18–22.

Lindh, M.V., Sjöstedt, J., Andersson, A.F., Baltar, F., Hugerth, L.W., Lundin, D., Muthusamy, S., Legrand, C., Pinhassi, J., 2015. Disentangling seasonal bacterioplankton population dynamics by high-frequency sampling: High-resolution temporal dynamics of marine bacteria. Environ. Microbiol. 17, 2459–2476. https://doi.org/10.1111/1462-2920.12720

Liu, C., Zhao, D., Ma, W., Guo, Y., Wang, A., Wang, Q., Lee, D.-J., 2016. Denitrifying sulfide removal process on high-salinity wastewaters in the presence of *Halomonas* sp. Appl. Microbiol. Biotechnol. 100, 1421–1426. https://doi.org/10.1007/s00253-015-7039-6

Liu, W., Xie, W., Zhao, Q., Zhu, K., Yu, R., 2014. Spatial distribution and ecological stoichiometry characteristics of carbon, nitrogen and phosphorus in soil in *Phragmites australis* tidal flat of Jiaozhou Bay. Wetl. Sci. https://doi.org/10.13248/j.cnki.wetlandsci.2014.03.014

Liu, X., Wu, J., Hong, Y., Jiao, L., Li, Y., Wang, L., Wang, Y., Chang, X., 2020. Nitrogen loss by *nirS*-type denitrifying bacterial communities in eutrophic coastal sediments. Int. Biodeterior. Biodegrad. 150, 104955. https://doi.org/10.1016/j.ibiod.2020.104955

Liu, Y.-X., Qin, Y., Chen, T., Lu, M., Qian, X., Guo, X., Bai, Y., 2021. A practical guide to amplicon and metagenomic analysis of microbiome data. Protein Cell 12, 315–330. https://doi.org/10.1007/s13238-020-00724-8

Louca, S., Parfrey, L.W., Doebeli, M., 2016. Decoupling function and taxonomy in the global ocean microbiome. Science 353, 1272–1277. https://doi.org/10.1126/science.aaf4507

Lu, M., Wang, X., Li, H., Jiao, J.J., Luo, X., Luo, M., Yu, S., Xiao, K., Li, X., Qiu, W., Zheng, C., 2022. Microbial community assembly and co-occurrence relationship in sediments of the river-dominated estuary and the adjacent shelf in the wet season. Environ. Pollut. 308, 119572. https://doi.org/10.1016/j.envpol.2022.119572

Luo, G., Fotidis, I.A., Angelidaki, I., 2016. Comparative analysis of taxonomic, functional, and metabolic patterns of microbiomes from 14 full-scale biogas reactors by metagenomic sequencing and radioisotopic analysis. Biotechnol. Biofuels 9, 51. https://doi.org/10.1186/s13068-016-0465-6

Lv, X., Ma, B., Yu, J., Chang, S.X., Xu, J., Li, Y., Wang, G., Han, G., Bo, G., Chu, X., 2016. Bacterial community structure and function shift along a successional series of tidal flats in the Yellow River Delta. Sci. Rep. 6, 36550. https://doi.org/10.1038/srep36550

Lyu, F., Han, F., Ge, C., Mao, W., Chen, L., Hu, H., Chen, G., Lang, Q., Fang, C., 2023. OmicStudio: A composable bioinformatics cloud platform with real-time feedback that can generate high-quality graphs for publication. iMeta 2. https://doi.org/10.1002/imt2.85

Lyu, X., Zhao, C., Xia, C., Qiao, F., 2010. Numerical study of water exchange in the Jiaozhou Bay and the tidal residual currents near the bay mouth. Acta Oceanol. Sin. 32, 20–30.

Magoč, T., Salzberg, S.L., 2011. FLASH: fast length adjustment of short reads to improve genome assemblies. Bioinformatics 27, 2957–2963. https://doi.org/10.1093/bioinformatics/btr507

Mori, F., Umezawa, Y., Kondo, R., Nishihara, G.N., Wada, M., 2021. Potential oxygen consumption and community composition of sediment bacteria in a seasonally hypoxic enclosed bay. PeerJ 9, e11836. https://doi.org/10.7717/peerj.11836

Morriën, E., Hannula, S.E., Snoek, L.B., Helmsing, N.R., Zweers, H., de Hollander, M., Soto, R.L., Bouffaud, M.-L., Buée, M., Dimmers, W., Duyts, H., Geisen, S., Girlanda, M., Griffiths, R.I., Jørgensen, H.-B., Jensen, J., Plassart, P., Redecker, D., Schmelz, R.M., Schmidt, O., Thomson, B.C., Tisserant, E., Uroz, S., Winding, A., Bailey, M.J., Bonkowski, M., Faber, J.H., Martin, F., Lemanceau, P., de Boer, W., van Veen, J.A., van der Putten, W.H., 2017. Soil networks become more connected and take up more carbon as nature restoration progresses. Nat. Commun. 8, 14349. https://doi.org/10.1038/ncomms14349

Mougi, A., Kondoh, M., 2012. Diversity of interaction types and ecological community stability. Science 337, 349–351. https://doi.org/10.1126/science.1220529

Narihiro, T., Nobu, M.K., Kim, N.-K., Kamagata, Y., Liu, W.-T., 2015. The nexus of syntrophy-associated microbiota in anaerobic digestion revealed by long-term enrichment and community survey: Microbial community of syntrophic enrichment cultures. Environ. Microbiol. 17, 1707–1720. https://doi.org/10.1111/1462-2920.12616

Nocker, A., Lepo, J.E., Martin, L.L., Snyder, R.A., 2007. Response of estuarine biofilm microbial community development to changes in dissolved oxygen and nutrient concentrations. Microb. Ecol. 54, 532–542. https://doi.org/10.1007/s00248-007-9236-z

Offret, C., Gauthier, O., Despréaux, G., Bidault, A., Corporeau, C., Miner, P., Petton, B., Pernet, F., Fabioux, C., Paillard, C., Le Blay, G., 2023. Microbiota of the digestive glands and extrapallial fluids of clams evolve differently over time depending on the intertidal position. Microb. Ecol. 85, 288–297. https://doi.org/10.1007/s00248-022-01959-0

Oksanen, J., 2010. Vegan: community ecology package.

Oyetibo, G.O., Ige, O.O., Obinani, P.K., Amund, O.O., 2021. Ecological risk potentials of petroleum hydrocarbons and heavy metals shape the bacterial communities of marine hydrosphere at Atlantic Ocean, Atlas Cove, Nigeria. J. Environ. Manage. 289, 112563. https://doi.org/10.1016/j.jenvman.2021.112563

Parks, D.H., Tyson, G.W., Hugenholtz, P., Beiko, R.G., 2014. STAMP: statistical analysis of taxonomic and functional profiles. Bioinformatics 30, 3123–3124. https://doi.org/10.1093/bioinformatics/btu494

Paterson, D.M., 1989. Short-term changes in the erodibility of intertidal cohesive sediments related to the migratory behavior of epipelic diatoms. Limnol. Oceanogr. 34, 223–234. https://doi.org/10.4319/lo.1989.34.1.0223

Pearce, D., Hodgson, D., Thorne, M., Burns, G., Cockell, C., 2013. Preliminary analysis of life within a former subglacial lake sediment in Antarctica. Diversity 5, 680–702. https://doi.org/10.3390/d5030680

Precht, E., Franke, U., Polerecky, L., Huettel, M., 2004. Oxygen dynamics in permeable sediments with wave-driven pore water exchange. Limnol. Oceanogr. 49, 693–705. https://doi.org/10.4319/lo.2004.49.3.0693

Qian, P.-Y., Lau, S.C.K., Dahms, H.-U., Dobretsov, S., Harder, T., 2007. Marine biofilms as mediators of colonization by marine macroorganisms: implications for antifouling and aquaculture. Mar. Biotechnol. 9, 399–410. https://doi.org/10.1007/s10126-007-9001-9

Regnell, O., Watras, Carl.J., 2019. Microbial mercury methylation in aquatic environments: a critical review of published field and laboratory studies. Environ. Sci. Technol. 53, 4–19. https://doi.org/10.1021/acs.est.8b02709

Reji, L., Tolar, B.B., Chavez, F.P., Francis, C.A., 2020. Depth-differentiation and seasonality of planktonic microbial assemblages in the Monterey Bay upwelling system. Front. Microbiol. 11, 1075. https://doi.org/10.3389/fmicb.2020.01075

Rickard, A.H., McBain, A.J., Stead, A.T., Gilbert, P., 2004. Shear rate moderates community diversity in freshwater biofilms. Appl. Environ. Microbiol. 70, 7426–7435. https://doi.org/10.1128/AEM.70.12.7426-7435.2004

Roggatz, C.C., Parsons, D.R., 2022. Potential climate change impacts on the abiotic degradation of acyl-homoserine lactones in the fluctuating conditions of marine biofilms. Front. Mar. Sci. 9, 882428. https://doi.org/10.3389/fmars.2022.882428

Saad, S., Bhatnagar, S., Tegetmeyer, H.E., Geelhoed, J.S., Strous, M., Ruff, S.E., 2017. Transient exposure to oxygen or nitrate reveals ecophysiology of fermentative and sulfate-reducing benthic microbial populations: Ecophysiology of benthic microbial populations. Environ. Microbiol. 19, 4866–4881. https://doi.org/10.1111/1462-2920.13895

Shang, H., Xi, M., Li, Y., Kong, F., Wang, S., 2018. Evaluation of changes in the ecosystem services of Jiaozhou Bay coastal wetland. Acta Ecol. Sin. 38, 421–431. https://doi.org/10.5846/stxb201608301763

Suh, S.-S., Park, M., Hwang, J., Kil, E.-J., Jung, S.W., Lee, S., Lee, T.-K., 2015. Seasonal dynamics of marine microbial community in the South Sea of Korea. PLOS ONE 10, e0131633. https://doi.org/10.1371/journal.pone.0131633

Sun, J., Wei, L., Yin, R., Jiang, F., Shang, C., 2020. Microbial iron reduction enhances in-situ control of biogenic hydrogen sulfide by FeOOH granules in sediments of polluted urban waters. Water Res. 171, 115453. https://doi.org/10.1016/j.watres.2019.115453

Sun, Q., Song, J., Li, X., Yuan, H., Xing, J., 2021. Spatial variations of bacterial community composition in sediments of the Jiaozhou Bay, China. J. Oceanol. Limnol. 39, 865–879. https://doi.org/10.1007/s00343-020-0127-1

Tavares, T.C.L., Bezerra, W.M., Normando, L.R.O., Rosado, A.S., Melo, V.M.M., 2021. Brazilian semi-arid mangroves-associated microbiome as pools of richness and complexity in a changing world. Front. Microbiol. 12, 715991. https://doi.org/10.3389/fmicb.2021.715991

Taylor, H.B., Kurtz, H.D., 2020. Microbial community structure shows differing levels of temporal stability in intertidal beach sands of the grand strand region of South Carolina. PLOS ONE 15, e0229387. https://doi.org/10.1371/journal.pone.0229387

Tebbe, D.A., Geihser, S., Wemheuer, B., Daniel, R., Schäfer, H., Engelen, B., 2022. Seasonal and zonal succession of bacterial communities in North Sea salt marsh sediments. Microorganisms 10, 859. https://doi.org/10.3390/microorganisms10050859

Thomas, F.A., Sinha, R.K., Krishnan, K.P., 2020. Bacterial community structure of a glacio-marine system in the Arctic (Ny-Ålesund, Svalbard). Sci. Total Environ. 718, 135264. https://doi.org/10.1016/j.scitotenv.2019.135264

Thomson, T., Ellis, J.I., Fusi, M., Prinz, N., Bennett-Smith, M.F., Aylagas, E., Carvalho, S., Jones, B.H., 2022. The right place at the right time: Seasonal variation of bacterial communities in arid Avicennia marina soils in the Red Sea is specific to its position in the intertidal. Front. Ecol. Evol. 10, 845611. https://doi.org/10.3389/fevo.2022.845611

van de Kamp, J., Hook, S.E., Williams, A., Tanner, J.E., Bodrossy, L., 2019. Baseline characterization of aerobic hydrocarbon degrading microbial communities in deep-sea sediments of the Great Australian Bight, Australia. Environ. Microbiol. 21, 1782–1797. https://doi.org/10.1111/1462-2920.14559

Wahl, M., Goecke, F., Labes, A., Dobretsov, S., Weinberger, F., 2012. The second skin: ecological role of epibiotic biofilms on marine organisms. Front. Microbiol. 3. https://doi.org/10.3389/fmicb.2012.00292

Wang, H., Zhang, C., Chen, F., Kan, J., 2020. Spatial and temporal variations of bacterioplankton in the Chesapeake Bay: A re-examination with high-throughput sequencing analysis. Limnol. Oceanogr. 65, 3032–3045. https://doi.org/10.1002/lno.11572

Wang, Y., Zhang, C., Qi, L., Jia, X., Lu, W., 2016. Diversity and antimicrobial activities of cultivable bacteria isolated from Jiaozhou Bay. Acta Microbiol. Sin. https://doi.org/10.13343/j.cnki.wsxb.20160132

Wang, Z., Gao, S., Sun, Z., Zhang, Y., Zhang, X., 2022. Study on the effects of rainfall and ecological water supplement on the pollutant transport in Licun River. Trans. Oceanol. Limnol. https://doi.org/10.13984/j.cnki.cn37-1141.2022.03.012

Wei, W., Isobe, K., Nishizawa, T., Zhu, L., Shiratori, Y., Ohte, N., Koba, K., Otsuka, S., Senoo, K., 2015. Higher diversity and abundance of denitrifying microorganisms in environments than considered previously. ISME J. 9, 1954–1965. https://doi.org/10.1038/ismej.2015.9

Wei, Z., Liu, Y., Feng, K., Li, S., Wang, S., Jin, D., Zhang, Y., Chen, H., Yin, H., Xu, M., Deng, Y., 2018. The divergence between fungal and bacterial communities in seasonal and spatial variations of wastewater treatment plants. Sci. Total Environ. 628–629, 969–978. https://doi.org/10.1016/j.scitotenv.2018.02.003

Whitehouse, R., 2000. Dynamics of estuarine muds: A manual for practical applications. Thomas Telford.

Wu, D., Huang, X.-H., Sun, J.-Z., Graham, D.W., Xie, B., 2017. Antibiotic resistance genes and associated microbial community conditions in aging landfill systems. Environ. Sci. Technol. 51, 12859–12867. https://doi.org/10.1021/acs.est.7b03797

Wu, J., Hong, Y., Ye, J., Li, Yiben, Liu, X., Jiao, L., Li, T., Li, Yuwei, Bin, L., Wang, Y., 2019. Diversity of anammox bacteria and contribution to the nitrogen loss in surface sediment. Int. Biodeterior. Biodegrad. 142, 227–234. https://doi.org/10.1016/j.ibiod.2019.05.018

Wyness, A.J., Paterson, D.M., Rimmer, J.E.V., Defew, E.C., Stutter, M.I., Avery, L.M., 2019. Assessing risk of *E. coli* resuspension from intertidal estuarine sediments: implications for water quality. Int. J. Environ. Res. Public. Health 16, 3255. https://doi.org/10.3390/ijerph16183255

Yi, J., Lo, L.S.H., Cheng, J., 2020. Dynamics of microbial community structure and ecological functions in estuarine intertidal sediments. Front. Mar. Sci. 7, 585970. https://doi.org/10.3389/fmars.2020.585970

Zhang, J., Zhang, N., Liu, Y.-X., Zhang, X., Hu, B., Qin, Y., Xu, H., Wang, H., Guo, X., Qian, J., Wang, W., Zhang, P., Jin, T., Chu, C., Bai, Y., 2018. Root microbiota shift in rice correlates with resident time in the field and developmental stage. Sci. China Life Sci. 61, 613–621. https://doi.org/10.1007/s11427-018-9284-4

Zhang, W., Tang, J., Liang, B., 2017. Numerical study on the influence of the forebay reclamation on pollutant transport in the Jiaozhou bay. Mar. Environ. Sci. https://doi.org/10.13634/j.cnki.mes.2017.01.005

Zhang, X., Hu, B.X., Ren, H., Zhang, J., 2018. Composition and functional diversity of microbial community across a mangrove-inhabited mudflat as revealed by 16S rDNA gene sequences. Sci. Total Environ. 633, 518–528. https://doi.org/10.1016/j.scitotenv.2018.03.158

Zhou, Z., Meng, H., Liu, Y., Gu, J.-D., Li, M., 2017. Stratified bacterial and archaeal community in mangrove and intertidal wetland mudflats revealed by high throughput 16S rRNA gene sequencing. Front. Microbiol. 8, 2148. https://doi.org/10.3389/fmicb.2017.02148

Zhuang, L., Tang, Z., Ma, J., Yu, Z., Wang, Y., Tang, J., 2019. Enhanced anaerobic biodegradation of benzoate under sulfate-reducing conditions with conductive iron-oxides in sediment of Pearl River Estuary. Front. Microbiol. 10, 374. https://doi.org/10.3389/fmicb.2019.00374

Zinke, L.A., Mullis, M.M., Bird, J.T., Marshall, I.P.G., Jørgensen, B.B., Lloyd, K.G., Amend, J.P., Kiel Reese, B., 2017. Thriving or surviving? Evaluating active microbial guilds in Baltic Sea sediment: Baltic Sea sediment metatranscriptomes. Environ. Microbiol. Rep. 9, 528–536. https://doi.org/10.1111/1758-2229.12578

