## Supplements for "Spatio-temporal variation of bacterial community structure in two intertidal sediment types of Jiaozhou Bay"

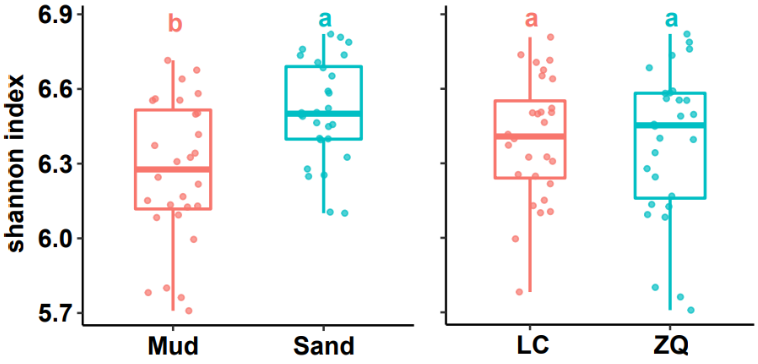


Fig. S1 Shannon index (TukeyHSD test) of samples by sediment type and by station

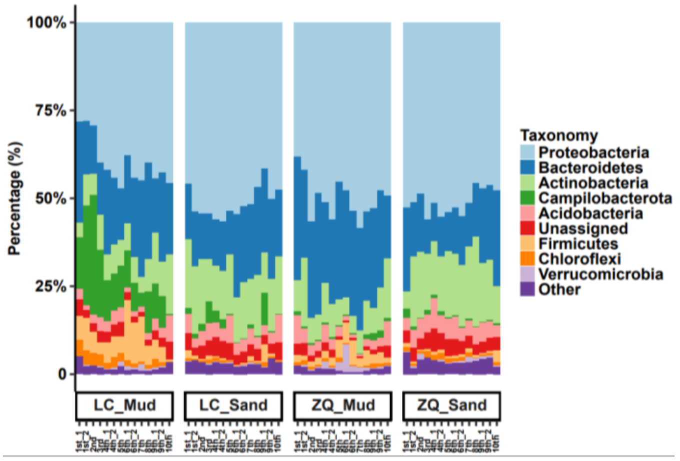


Fig. S2 Relative abundance at major phylum levels

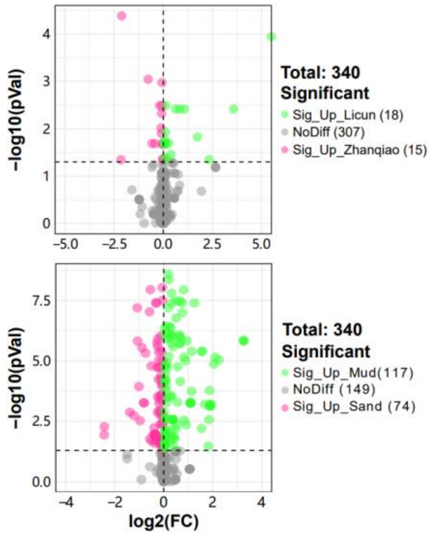


Fig. S3 KEGG_L3 volcanogram analysis across stations and substrates.

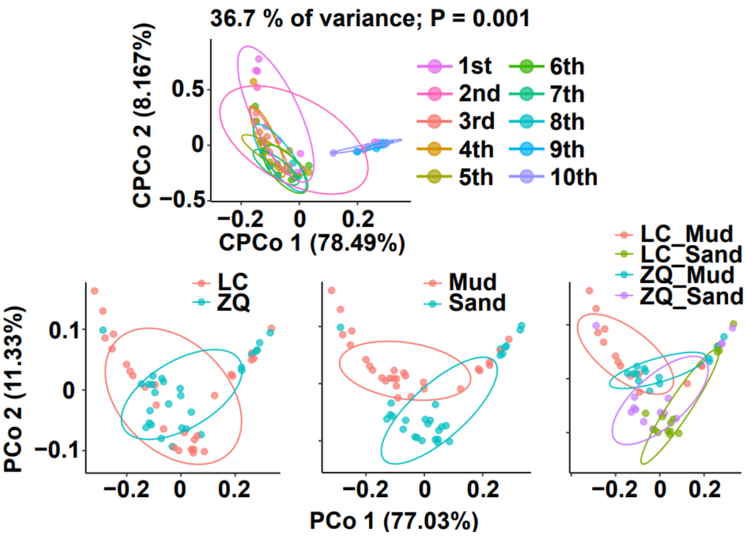


Fig. S4 KO CPCoA by month and KO PCoA (Bray_Curtis) of samples from different stations and substrates

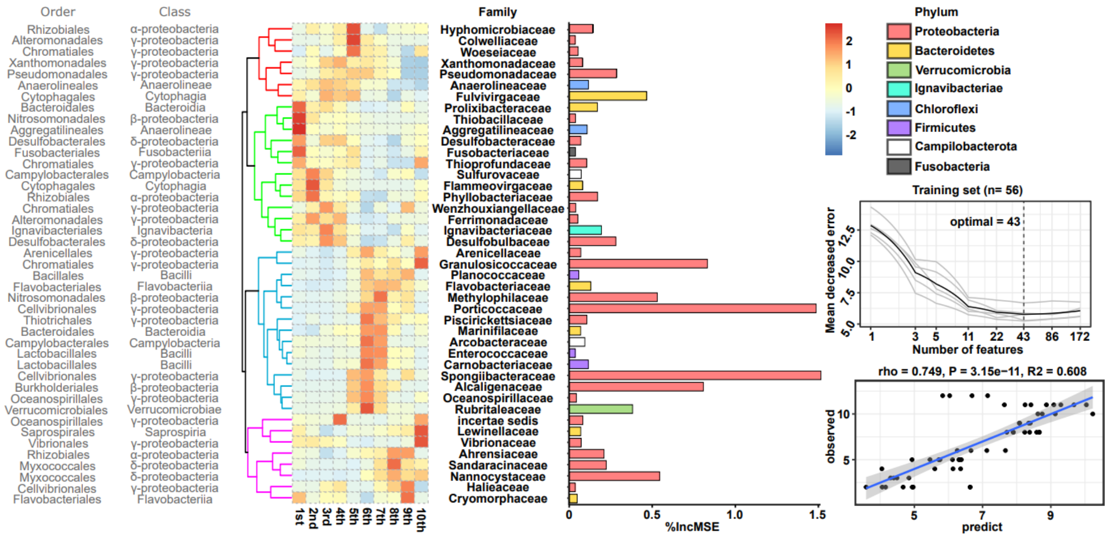


Fig. S5 Plots of random forest predictions at the family level of temporal correlation features abundance, importance, cross-validation and temporal prediction fit.
